# Auditory attention detection in cocktail-party: A microstate study

**DOI:** 10.1101/2023.09.27.559867

**Authors:** Hossein Roushan, Sahar Bakhshalipour Gavgani, Masoud Geravanchizadeh

**Affiliations:** University of Tabriz

**Keywords:** selective auditory attention, EEG microstates, dichotic listening, HRTF conditions, cocktail-party problem

## Abstract

Healthy human brain can easily attend to one auditory stimulus and filter out the other stimuli, described in the terms of “selective auditory attention” or referred to as the cocktail party phenomenon. This paper proposes a new real-time mapping-based selective auditory attention detection (SAAD) system using only EEG signals. In the proposed system, several quasi-stable maps, called EEG microstates, are extracted. After the optimization of the microstates topographies, the temporal dynamics of the microstates are used for the classification of auditory attention direction. The evaluation results obtained in different listening conditions (i.e., dichotic and spatial hearing) show that the use of an integrated set of 6 microstate topographies best describes the data and provides better classification performance. Due to the nature of EEG microstates in reflecting the fundamentals of brain information processing, this study provides an appropriate way of reaching the underlying mechanism of selective attention. The proposed system based on microstates can effectively detect the attention direction in dynamic scenarios, where the listener’s attention might be switched in seconds, making the model suitable for real-time applications. Furthermore, the proposed auditory attention detection system is advantageous in the sense that the detection of attention is performed without using speech signals.

---

One of the distinctive abilities of the human is to focus on one speaker in a complex auditory scene comprising multiple sound sources. This phenomenon which is widely known as the cocktail party effect was first established by Cherry (Cherry, 1953) and has been studied since then to understand the sensory solutions to the cocktail party problem (Bee & Micheyl, 2008; Bregman, 1994; Carlyon, 2004; McDermott, 2009; Shamma & Micheyl, 2010). For hearing-impaired listeners, it is challenging to separate the target signal (i.e., the talker of interest) from interferences (i.e., all other talkers) in a cocktail party scene with multiple people talking simultaneously. Accordingly, identifying the target in hearing devices is crucial to improve the listening ability in complex acoustic environments (Carhart & Tillman, 1970; Glyde et al., 2013; Kidd Jr et al., 2015; Plomp, 1994; Zeng et al., 2008). Hearing devices cannot segregate and enhance the desired signal without knowing which talker the listener is attending to. The selective auditory attention detection (SAAD) approach proposes a solution, which detects the attention direction and helps to isolate and enhance the listener’s attended signal (Das et al., 2020; Geravanchizadeh & Zakeri, 2021).

To identify the target speaker, various SAAD methods have been proposed based on two groups of approaches: (1) using stimulus-response modeling, and (2) informative features from brain response signals. The first group of SAAD approaches begins with the realization that the amplitude envelope of the attended stream modulates the neural responses (Ding & Simon, 2012; Holmes et al., 2018). These findings have led to one of the most used methods for SAAD, named temporal response functions (TRFs), which is based on the design of mapping decoders to reconstruct the amplitude envelopes of the stimuli (Mesgarani & Chang, 2012). Afterward, successful progress was achieved by making many modifications to this basic mapping approach. The large training data for the mapping technique and the number of electrodes used in (O’Sullivan et al., 2015) were reevaluated in (Mirkovic et al., 2015) in which the required electrodes were reduced while maintaining the performance at an acceptable value. Since robust detection performance in this approach is highly dependent on various factors, such as the subject, multiple trials for training, and the temporal resolution, reducing training data causes lower accuracies both for the subject-dependent and subject-independent cases. The need for a large amount of data makes the TRF-based approaches not to be well-suited for emerging real-time applications such as smart hearing aid designs or brain-computer interfaces. To establish an algorithmic pipeline for real-time decoding, state-space modeling was developed to track the attentional state of the listener (Akram et al., 2016; Miran et al., 2018). Moreover, various regularization methods were applied to the mapping procedure to maintain the improved performance of the TRF technique (Wong et al., 2018). Another attempt to decode attention from short segments of EEG signals (10 seconds or less in duration) was made by extracting the brainstem response from multi-channel scalp recordings (Etard et al., 2019), where, it is demonstrated that the achieved accuracy of SAAD is similar for both subject-dependent and subject-independent training. Here, the attention decoding using only three electrodes achieves an average accuracy of 69% and 72%, with 16 seconds and 32 seconds of data, respectively. To investigate the influence of head-related filtering and ear-specific decoding, a spatio-temporal decoder is trained and used to perform SAAD (Das et al., 2016). In another study using both EEG recordings and speech streams, the auditory attention was decoded without a mapping procedure by extracting the features of EEG signals and auditory stimuli (Haghighi et al., 2018; Horton et al., 2014). The SAAD system in (Ciccarelli et al., 2019) proposes the end-to-end classification system with integrated similarity computation between EEG signals and audio envelopes. In another attempt, a dynamic SAAD system was introduced in which the recurrent neural network (RNN) and reinforcement learning (RL) methods were incorporated into the detection model (Geravanchizadeh & Roushan, 2021).

In the second group of methods for SAAD, the attention direction of listeners is detected by extracting appropriate informative features from the response signals and using different machine-learning approaches as classifiers. One approach in this group was given in (Thorpe et al., 2012), where the direction of the attention was detected by the classification of the EEG spectra with a naïve Bayes classifier plus feedback from subjects to improve the classification performance. In another study, it was shown that the use of entropy measures in EEG signals could be used to detect the physiological and cognitive states of the human brain (Lu et al., 2018), where, the subjects’ auditory attention from single-trial EEG (i.e., 60 seconds of EEG) was determined by the use of entropy measures. The cocktail party experiment paradigm in (Tian et al., 2018) was closer to the real-world condition with 89.7% accuracy, in which a feature extraction method, called rhythm entropy was adopted to classify the error-related auditory attention. Considering attention as a higher brain function, another study employed the connectivity analysis and graph-based features of the acquired connections to detect the attention direction with a gain of 93.5% accuracy in 60 seconds of EEG recordings (Geravanchizadeh & Gavgani, 2020).

As it is widely understood, the most used sensing modality for SAAD is EEG which is an inexpensive, yet powerful method for measuring brain electrophysiology with high temporal and fairly good spatial resolution. A distinct powerful approach to studying EEG signals is EEG microstates analysis which represents spatio-temporal dynamics of large-scale data and is used in a wide range of clinical and cognitive brain studies (D’Croz-Baron et al., 2019; Koenig et al., 2002; Michel & Koenig, 2018; Milz et al., 2016). EEG microstates analysis gives fundamental observation of human brain functioning and is used to investigate brain electric activity at a millisecond resolution by considering the multichannel EEG recording as a series of quasi-stable “microstates” (Koenig et al., 2002). It is suggested that the fundamental aspects of self-consciousness can be explored using EEG microstates (Bréchet & Michel, 2022). Task-related microstate analysis is a method for studying brain function and dysfunction that has some advantages over resting-state microstate analysis (Chen et al., 2021). One of the main advantages of task-related microstate analysis is that it can track changes in brain activity over time and how they are influenced by the task or stimulus. In contrast, resting-state microstate analysis can only measure the stable features of brain activity. This means that task-related microstate analysis is a more effective and informative method than resting-state microstate analysis for understanding and assessing brain network dynamics on a millisecond timescale.

In this study, a novel SAAD system is proposed by exploiting brain microstates of individuals who are listening to speech stimuli in both the dichotic listening scene and the spatially filtered configuration using Head-Related Transfer Functions (HRTFs). HRTF-filtered datasets are more realistic, in the sense the stimuli are propagated to both ears at different spatial angles. Here, the final goal is to find an optimized integrated microstates topography set which is used to detect auditory attention by considering EEG microstates dynamics with 2 seconds of EEG recordings. Accordingly, first, various sets of microstate maps are evaluated and the most describing microstates map set is selected via meta-criterion. Then, the corresponding dynamics are fed to a neural network (NN) classifier to detect the direction of selective auditory attention.

The organization of the paper is as follows. First, in “Materials and methods”, the methodology including the data description and the proposed microstate-based SAAD is explained. The details of EEG microstates determination and feature extraction method are presented in this section. Next, the section “Experiments and results” covers the experiments and evaluation results. Finally, in “Discussion and conclusion”, we conclude the paper with a summary of our findings, the discussions of their implications, and some recommendations for future studies.

## Materials and methods

### Data description and preprocessing

To ensure the validity of the obtained results in a more realistic scenario, the publicly available auditory attention detection dataset from ExpORL, Dept.

Neurosciences, KULeuven containing two listening conditions (i.e., Dichotic and HRTF) is used (Das et al., 2019). This dataset includes 16 normal-hearing subjects who participated in the experiment, listening to stimuli from four Dutch short stories narrated by different male speakers. The dichotic condition is where the audio streams are administered to separate channels of the insert phones at equal intensities without any filtering. The HRTF condition instead, is where the two audio streams are filtered by HRTFs, simulating an auditory environment in which, each speaker perceives the stimuli at 90 degrees to the left and right of the speaker. The subjects were asked to focus only on one of the streams. During the experiment, a total of 72 minutes of EEG signals in 20 trials for each subject, half in dichotic condition and half in HRTF condition, is recorded by the BioSemi system with 64 electrodes at 8196 Hz sampling rate. To make the dataset ready, the EEG recordings are re-referenced to the average of all scalp channels and downsampled to 128 Hz. Trials numbers 9 to 20 are extracted due to their equal size of time frames to use for the computation of the grand average. Then, all signals are segmented into 2-second epochs to compute microstates series.

### Proposed microstate-based SAAD

To identify the attention direction of subjects (i.e., attending to the left or right streams) with EEG microstates analysis, the artifact-free EEG signals are used as inputs for the proposed SAAD model. In the first stage, the preprocessed EEG recordings are used to compute the grand average (GA) and the global field power (GFP). Next, for the grand averages of trials 9 to 20, different numbers of microstate class topographies (here, from 1 to 20) are computed using the Topographic Atomize and Agglomerate Hierarchical Clustering (T-AAHC) algorithm. These topographies are then compared using global explained variance (GEV) and meta-criterion to obtain the optimal number of microstate topographies (Custo et al., 2017). After the determination of the optimal topography, the obtained maps are fitted to all the data. In the next stage, the dynamics of EEG microstates, including the duration and coverage of each microstates class are extracted. Finally, the computed features are fed into a NN classifier to detect the attention direction.

#### Microstate analysis

EEG studies on cognitive processes (e.g., behavior, thoughts, and emotions) demonstrate that global electrical brain activity can be classified into a finite number of quasi-stable topographies for transient periods, referred to as ‘EEG microstates’(Michel & Koenig, 2018). These repeating microstates and their sequences are strongly related to the content of the mind.

The microstates analysis begins by computing the grand average (GA) of the preprocessed EEGs to determine the microstate classes using the T-AAHC algorithm. The T-AAHC is a bottom-up approach that starts with all EEG time frames being a cluster map. Then, it picks the “worst” cluster of all currently available ones to atomize it (i.e., split it into individual maps), and to re-distribute these maps to the segments they fit most. It then merges the two closest clusters (of the current remaining ones) into a new one, repeatedly. All the segmentations from the number of time frames down to 1 are obtained in this way. This method has many advantages over the more conventional K-Means method such as running much faster and giving the same results in different iterations as it does not depend on the initialization. After the determination of microstate classes, the analysis continues with computing GFPs and their peaks (Koenig et al., 2002). The GFP indicates the strength of the electric field over the brain and is technically the standard deviation of the temporal potential values at all electrodes. It is used for measuring the global brain response to an event or characterizing the fast modifications in brain activity (Lehmann & Skrandies, 1980). Next, the EEG time points at the neighborhood of the GFP peaks are fed into a second T-AAHC algorithm to assign them to a microstate class if its correlation with the cluster map is above 0.5 (Michel et al., 2009).

In more conventional studies, brain activity at the resting state is defined with 4 number of microstates (Brodbeck et al., 2012; Lehmann et al., 2005; Pascual-Marqui et al., 1995), while the number of microstates in task mode is usually unknown (Kindler et al., 2011; Koenig et al., 1999). To represent brain activity for a specific dataset, there is a need to determine an optimal set of topographies. This is necessary for capturing the most describing mapping and informative features of the data and avoiding over or underfitting (Li et al., 2007; Michel & Koenig, 2018). The optimal number of microstates class maps can be determined by an optimization criterion named the meta-criterion (Bréchet et al., 2020). In this criterion, the optimal number of microstate topographies is obtained by the median of all optimal numbers of clusters (Bréchet et al., 2019; Custo et al., 2017) across 7 different criteria including, Gamma, Silhouettes, Davies and Bouldin, Point-Biserial, Dunn, Krzanowski-Lai Index, and Cross-Validation (Charrad et al., 2014; Krzanowski & Lai, 1988; Milligan & Cooper, 1985; Pascual-Marqui et al., 1995). The meta-criterion gives a more reliable result from all criteria, where each single criterion uses a distinct metric to evaluate the “quality” of a single segmentation. After finding the optimal number of microstates, GEV is used to measure how well the microstate sequence approximates the underlying EEG data set (Murray et al., 2008).

In this study, the analysis was performed using the Cartool software developed by Denis Brunet (cartoolcommunity.unige.ch).

#### Calculation of Microstate Parameters

Different parameters of EEG microstates explain the characteristics of neural activity (Fu et al., 2014; Khanna et al., 2015). Three temporal dynamics parameters are used in this study, including coverage, duration, and occurrence of each microstate map. The fraction of the total recording time that a microstate is dominant is called the coverage. The duration of each microstate map is related to the temporal stability and is defined by the average duration that a certain microstate remains stable. The frequency of occurrence for each microstate topography, independent of its duration, defines the third parameter.

### Experiments and results

In this paper, we apply and evaluate the proposed microstate approach in selective auditory attention detection tasks. Accordingly, the first step after preprocessing the EEG data is to find the optimal number of microstate topographies that best represent the brain activity during the listening task. In this regard, EEG microstate topographies are calculated with different numbers of microstate classes, ranging from 1 to 20 maps in three different experiments; finding A) dico, B) hrtf, and C) intg maps. These EEG microstates are explored by a statistical method called meta-criterion, which evaluates the goodness of fit and the complexity of the microstate models, to determine the optimal number of maps that capture the most relevant information with the least redundancy. In an additional experiment, the conventional 4-class microstates map set (i.e., conv maps) is used as a baseline for comparison with newly found topographies.

### Computing optimal maps

#### dico maps

Dichotic listening condition is a kind of auditory test that involves playing different sounds to each ear simultaneously, usually using headphones, which can be used to investigate selective attention and the lateralization of brain function within the auditory system. The EEG recording in this experiment covers two types of scenarios: listening to the left stream and listening to the right stream. This experiment aims to discover a set of maps (i.e., dico maps) that can be used to detect auditory attention for listeners in a dichotic setting. Based on the meta-criterion, 4 microstate topographies give the best explanation for the EEG data of dichotic listeners.

#### B. hrtf maps

The HRTF condition simulates the spatial information of sound sources located in different positions in space. This condition tests how listeners can focus their attention on a specific sound direction in a complex acoustic scene. In this experiment, the EEG recordings of those who paid attention to the left or right stream in the HRTF condition are used. The goal of this experiment is to find a set of maps (i.e., hrtf maps) that can help identify auditory attention for listeners in the more complicated listening condition. Using the meta-criterion, 6 microstate topographies provide the best account for the EEG data of listeners to spatially filtered streams.

#### C. intg maps

In this experiment, the EEG recordings of all four groups of listeners, including left-stream and right-stream listeners in both dichotic and HRTF conditions, are used. Here, using common neural activities underlying attention in both conditions, we aim to find an integrated set of maps (i.e., intg maps) that is applicable for detecting auditory attention, regardless of the listening scenario. According to the meta-criterion, 6 microstate topographies provide the best explanation for the EEG data of all classes of listeners.

Figure 1 shows microstate maps that are found for each experiment showing their spatial distribution and polarity.

**Figure 1.**
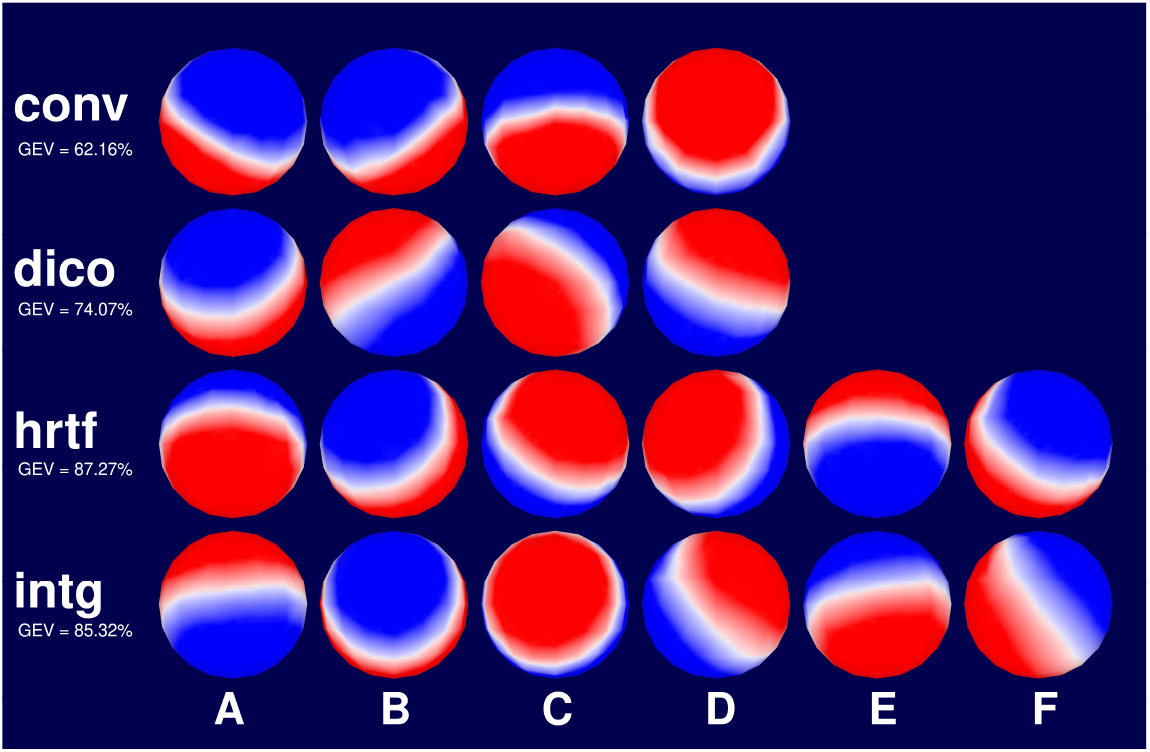
The optimal microstate topographies extracted in each experiment; from top to bottom: conv maps, dico maps, hrtf maps, and intg maps.

### Fitting maps

After finding the optimal microstate topographies that best fit the sample data, these topographies are assigned to the entire data set of EEG recordings and transform EEG signals into microstates segments. To make sure that these maps explain our data well, the GEV measure is employed. GEV quantifies how much of the variance in EEG data is accounted for by each microstate topography. The results show that the best set of maps that explain the most variance in our recordings is the ‘hrtf’ map set with GEV=87.27%. The next best-explaining topographies are ‘intg’ maps followed by ‘dico’ maps with GEV=85.32% and GEV=74.07%, respectively. The ‘conv’ map set shows the least explainability of the data variance with GEV=62.16% (see Fig. 1).

The pairwise spatial correlation among the optimal microstate topographies is calculated to further examine the quality of computed microstate maps. This is done to inspect the possibility of having maps similar to each other which makes one of them redundant. The spatial correlation is a measure of how much two maps overlap in their spatial distribution and polarity. Figure 2 shows the matrix of spatial correlations among maps in three different experiments.

**Figure 2.**
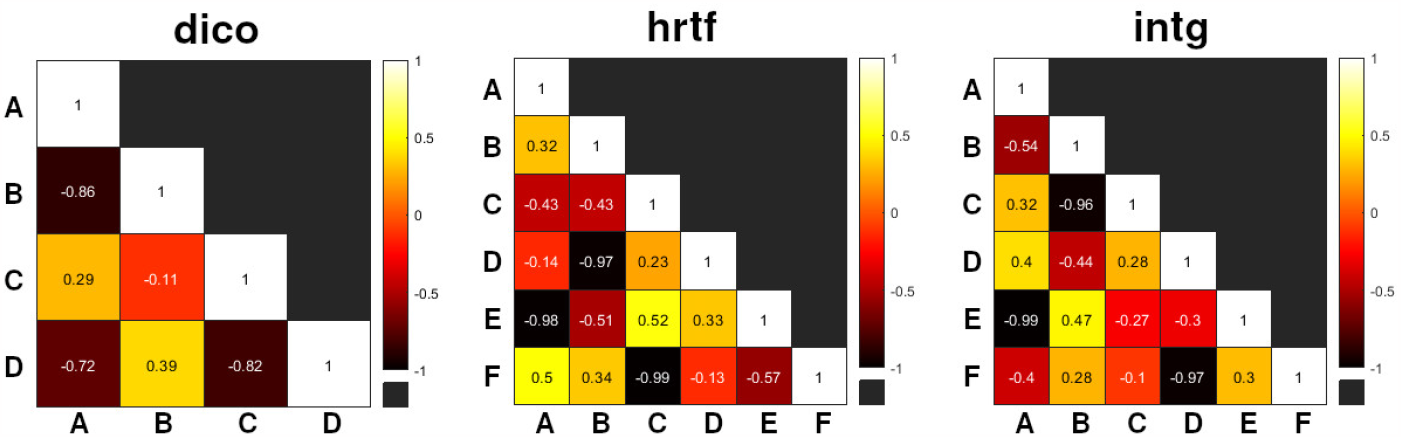
The pairwise spatial correlation between the optimal microstate maps (from left to right) for dico, hrtf, and intg maps.

### Temporal dynamics

The next step of the study is to extract features from EEG microstates segments to feed the classifier. To do this, three microstate parameters, known as temporal dynamics, are calculated using the segments of the EEG microstates: Coverage, Duration, and Occurrence. Coverage is the percentage of time that each microstate topography occupies in the microstates segment, while duration is the average time that each microstate topography lasts in the segment. Occurrence is the frequency of having a particular map, regardless of its duration. Figure 3 shows the average amount of temporal dynamics for each map in all experiments.

**Figure 3.**
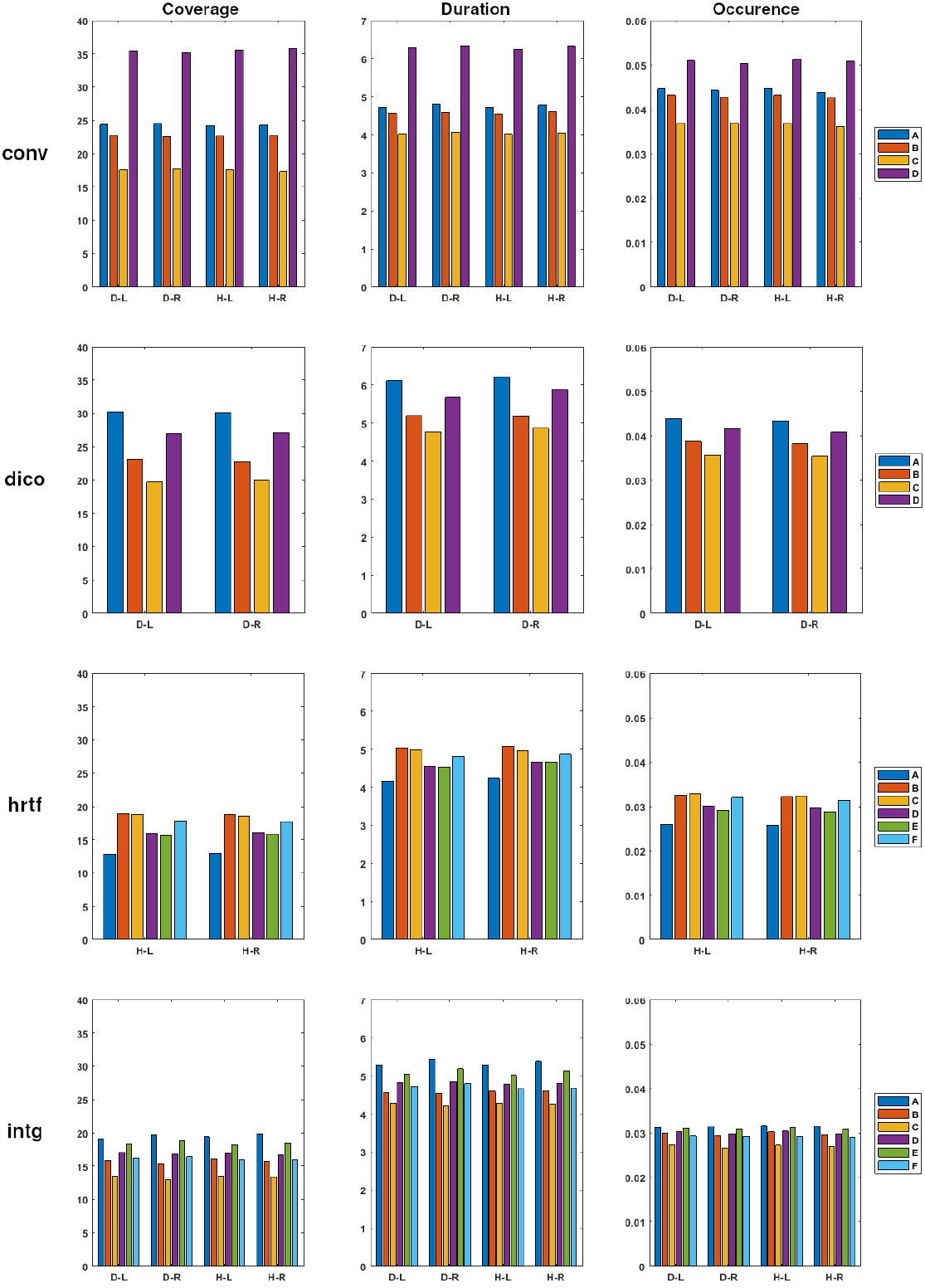
The average amount of temporal dynamics for each map in ‘conv’, ‘dico’, ‘hrtf’, and ‘intg’ experiments. Here, the labels D-L, D-R, H-L, and H-R, represent the data used in dichotic (‘D’) and HRTF (‘H’) conditions for the left (‘L’) and right (‘R’) attending cases.

### Classification

In the last step of experiments, the three parameters calculated in the previous stage are used as input features for the NN classifier to detect the direction of attention. The classifier is composed of a multi-layer perceptron neural network, with two hidden layers. The first hidden layer has 10 fully connected neurons, and the second hidden layer has 100 fully connected neurons. The activation function used in both hidden layers is a rectified linear unit (ReLU). In each run of the algorithms, 70% of the data is selected at random for the training of the NN classifier. The rest is taken for the test procedure. To avoid the overfitting issue in the detection of classes, a 5-fold cross-validation technique is employed. The results of the classification in different experiments are shown in Table 1. Here, the accuracy is measured as the percentage of correctly classified trials averaged over 10 runs shown with standard deviations.

**Table 1.** The results of proposed microstate-based SAAD in terms of Accuracy for different listening conditions and microstate map sets.

| Condition | Template | Accuracy (%) |
| --- | --- | --- |
| Dichotic | conv | $71.45 \pm 1.75$ |
| | dico | $73.48 \pm 1.92$ |
| | intg | $82.07 \pm 0.60$ |
| HRTF | conv | $73.46 \pm 1.41$ |
| | hrtf | $82.07 \pm 1.05$ |
| | intg | $81.59 \pm 0.90$ |

## Discussion and conclusion

Task-related microstate analysis is an effective and informative method compared to resting state microstate analysis for understanding and assessing brain network dynamics on a millisecond timescale. This study aims to introduce an integrated set of microstates topographies that best describes the brain activity in a selective auditory attention task and develop a new SAAD system based on the analysis of EEG microstates. The proposed system can track the temporal changes in the listener’s attention direction using the EEG signals recorded from the scalp of the listener without requiring speech stimuli, which are often used in traditional methods for auditory attention detection. The proposed method processes the EEG signals in different steps. Initially, it calculates the GFP, which is a measure of the overall strength of the EEG signals. Then, it determines the optimal number of clusters for different numbers of microstates. Here the optimal number of clusters is found by comparing the fit of different models using a measure called meta-criterion. Finally, it extracts the corresponding dynamics of EEG microstates, which describe how the microstates change over time. The system then uses the microstate temporal dynamic features as input for the NN classifier to detect the attention direction.

The proposed system has several advantages over the conventional methods for auditory attention detection. First, it relies solely on the brain activity potentials, which simplifies the real-world applications of SAAD. Second, the detection process is performed in short time segments (i.e., 2 seconds) with a significantly high accuracy. This means that the algorithm can track the attention direction in real-time. Third, it can be applied to various auditory conditions (i.e., dichotic and HRTF), which makes it more versatile and robust.

The idea of employing optimal map sets for the selective auditory attention detection task was tested in various listening conditions. Table 1 presents the accuracy of the proposed microstate-based SAAD method for different listening conditions and microstate map sets. In this table, the listening conditions are dichotic and HRTF and the microstate map sets are represented as *conv, dico, hrtf*, and *intg*, which are acquired in different experiments. The *conv* maps result in the lowest accuracies among all microstate templates. This was predictable since this map set could not cover the particular information associated with the auditory attention task. This has motivated the authors to explore and compute optimal map sets specifically for SAAD in different listening scenarios. The results indicate that the *hrtf* map set, which uses only data in the HRTF condition, performs significantly better classification than *conv* maps, while the optimal *dico* maps achieve relatively lower improvements. This suggests that the *hrtf* maps are more sensitive to the spatial cues provided by the HRTF stimuli. The *intg* map set, which integrates data from both dichotic and HRTF conditions, yields considerably high accuracy for both listening conditions. The high performance of this map set can be justified by the fact that it captures the most relevant EEG features for SAAD, regardless of the condition. These results are also supported by the computed GEV values for different optimal map sets shown in Figure 1. In other words, the templates which better explain the EEG recordings, result correspondingly in higher attention detection accuracies. In general, the classification performances achieved in the HRTF condition are higher than those in the dichotic condition which shows the importance of spatial auditory cues used in SAAD. The possible redundancy of extracted topographies is further investigated by calculating the pairwise correlations between optimal microstate maps (Fig. 2). In all experiments, it is revealed that spatial correlations between different maps are lower than 0.52, which results in a lower possibility of mislabeling two microstates. To probe the advantages of integrated topographies (i.e., *intg* maps) over conventional microstates (i.e., *conv* maps), the pairwise spatial correlations between these two map sets are calculated and shown in Figure 4. It is seen that maps *intg*-F, *intg*-B, *intg*-E, and *intg*-C have notably high correlations with *conv*-A, *conv*-B, *conv*-C, and *conv*-D, respectively, which means that they have common information. Maps *intg*-A and *intg*-D have low correlation values with *conv* maps which can be interpreted as the least similarity. As a result, in the case when only four conventional maps are employed, the information of microstates in these two topographies is missed. In other words, maps *intg*-A and *intg*-D represent distinct brain states that contain discriminating auditory attention information.

**Figure 4.**
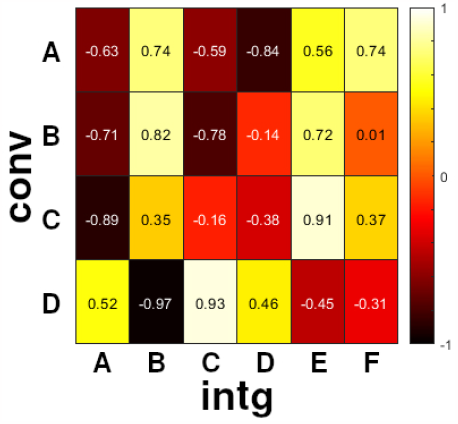
The pairwise spatial correlation between the conv maps and intg maps.

The current work evaluates the performance of the SAAD system in dichotic and HRTF scenarios. The assessment of the proposed system with more sound sources located at different spatial positions in a real noisy and reverberant acoustic environment seems to be an indispensable step toward the design of practical hearing aids. This step is going to be achieved in our future work. Furthermore, switching attention and detection time are important factors in SAAD, especially for applications that require real-time feedback or adaptation. Although the proposed SAAD performs with only 2 seconds of EEG recordings, which is comparatively faster than other methods, in future works, advanced dynamic detection methods should be incorporated in the proposed SAAD to detect and track the auditory attention switching in real environments.

## Acknowledgement

The Cartool software (cartoolcommunity.unige.ch) has been programmed by Denis Brunet, from the Functional Brain Mapping Laboratory (FBMLab), Geneva, Switzerland, and is supported by the Center for Biomedical Imaging (CIBM) of Geneva and Lausanne.

